# Programmable design of functional RNAs through multimodal generative learning

**DOI:** 10.64898/2026.10.14.759035

**Authors:** Jianbo Ma, Jianghui Zhu, Wenze Huang, Kunting Mu, Kaile Huang, Zhuoer Dong, Fengmin Guo, Xin Xie, Yanqiu Shao, Roland G. Huber, Yue Wan, Linqi Zhang, Qiong Wu, Qiangfeng Cliff Zhang

**Affiliations:** State Key Laboratory of Membrane Biology, Beijing Advanced Innovation Center for Structural Biology & Frontier Research Center for Biological Structure, School of Life Sciences, Tsinghua University; Beijing 100084, China; MOE Key Laboratory of Bioinformatics, Center for Life Sciences and Artificial Intelligence, Center for Synthetic and Systems Biology, School of Life Sciences, Tsinghua University; Beijing 100084, China; Tsinghua-Peking Center for Life Sciences; Beijing 100084, China; MOE Key Laboratory of Bioinformatics, State Key Laboratory of Green Biomanufacturing, Center for Synthetic and System Biology, School of Life Sciences, Tsinghua University; Beijing 100084, China; Comprehensive AIDS Research Center, Pandemic Research Alliance Unit, Center for Infection Biology, School of Basic Medical Sciences, Tsinghua University, Beijing 100084, China; Xungene Therapeutics; Hangzhou 310000, Zhejiang, China; Bioinformatics Institute, Agency for Science, Technology and Research, Matrix 07-01; Singapore 138671, Singapore; Genome Institute of Singapore, Agency for Science, Technology and Research, Genome 02-01; Singapore 138672, Singapore

## Abstract

RNA engineering is bottlenecked by the tight coupling among sequence, structure, and function, leaving the field reliant on heuristics or screening. We introduce RNAverse, a multimodal generative model that learns a conditionable design grammar from RNA sequences, experimental structural probing profiles, and functional annotations. Beyond setting a new state of the art across 16 predictive modeling tasks, RNAverse inverts into generative design across four distinct RNA classes through three regimes, requiring experimental testing of no more than 20 candidates per-task. Zero-shot in-painting generates snoRNAs that promote premature termination codon (PTC) readthrough, and chimeric aptazymes for ligand-controlled gene regulation, exceeding rational design or large-scale screens. In both settings, RNAverse outperforms family-adapted sequence-only baselines. Fine-tuned in-painting extends to an unseen functional class, yielding IGFBP3-binding aptamers with picomolar affinity that outperform SELEX winners. Iterative optimization engineers 5′ UTRs that outperform clinical benchmarks, enhancing protein production in cell-based assays for both the Zolgensma gene therapy and the BNT162b2 COVID-19 vaccine, and boosting antibody responses to BNT162b2 in mice. Together, these results establish multimodal generative learning as a route from predictive RNA modeling to programmable specification of functional RNAs.

## Introduction

RNA has emerged as a versatile substrate for medicine and synthetic biology: it can recognize specific RNA targets, sense metabolites, recruit protein machinery, catalyze reactions, and program protein expression^1–4^. These capabilities have already been realized in the clinic, from mRNA vaccines deployed at global scale to approved antisense and siRNA drugs, aptamers, and RNA-based gene therapies^5^. Yet engineering new functional RNAs remains dependent on rational heuristics^6^ or high-throughput screening (HTS) of large libraries^7–9^, both of which explore only a vanishing fraction of molecular space and are often tailored to individual functions. The challenge is one of specification: the relationship between RNA sequence and function is many-to-many: dissimilar sequences can preserve the same folded structure and function^10^, while nearly identical sequences can diverge through differences in molecular structure, regulation, and cellular milieu^11–13^. Programmable RNA engineering, *i.e.*, the design of sequences that, by construction, realize user-specified functions, requires traversing this map in reverse: a direction in which sequence alone provides little foothold.

Data-driven approaches have followed two recurring strategies that attempt to circumvent this obstacle. The first is family-specific generation: generative models are built for a single RNA family or functional class, either from evolutionary sequence covariation (*e.g.*, RfamGen^14^) or directly from sequence language models trained on a particular functional class (e.g., GEMORNA^15^, which autoregressively generates UTRs and coding sequences, along with approaches that exploit sequence likelihoods to propose stabilizing mutations, as demonstrated for improving rRNA thermostability^16^). These methods generate new family members effectively when training data are deep, but are confined to one family or functional class at a time and fail when family-specific data are sparse. The second strategy is predictor-guided search: a generative model is coupled with a task-specific predictor; candidate sequences are generated, scored for function, and iteratively refined toward better performance (*e.g.*, RNAGenesis^17^ and GARDN-SANDSTORM^18^). While this generate–predict–refine loop is powerful, it demands new data and a newly trained predictor for every task. RNA design thus remains a collection of bespoke pipelines, each rebuilt for every new family or task.

We reasoned that the missing ingredient is modality itself. At root, functional information never enters the generative model as an explicit conditioning signal: it is fixed by the training set in one strategy and delegated to an external predictor in the other. Indeed, functional annotations can provide the semantic layer that allows parameters to be shared across mechanistically distinct RNA classes and design intent to be specified at inference time, directly addressing the one-family-one-model limitation. As a complement, experimentally measured RNA structure, in the form of transcriptome-wide chemical probing at nucleotide-resolution^19,20^, provides a context-dependent view of RNA molecules precisely where structural state is measurable and functionally consequential. Together, these modalities create an opportunity to learn a shared design grammar in which sequence provides molecular syntax, functional annotations the semantics that enable cross-family generalization, and structural profiles additional experimentally grounded biophysical context.

Here we introduce RNAverse (RNA versatile embedding representation space for engineering), a multimodal generative model for specification-guided RNA design. RNAverse is trained in two stages: large-scale sequence-only pretraining on 540 billion nucleotides to capture broad regularities, followed by multimodal alignment with experimental structural profiles and functional annotations. RNAverse achieves leading performance across 16 predictive RNA modeling benchmarks among the evaluated models, with the clearest gains on tasks involving structural context, supporting the role of the added structural profiles in expanding its representational capacity. The decisive test, however, is whether this shared representation enables the design of functional RNAs across mechanistically distinct classes.

Our central result is RNA design by *zero-shot in-painting*: a single shared backbone designs functional RNAs from a functional specification alone, yielding high-potency PTC-readthrough snoRNAs and high-dynamic-range ligand-responsive aptazymes without task-specific retraining—and in both settings, RNAverse outperforms sequence-only baselines even when those baselines are fine-tuned on the corresponding families. This moves beyond both prior strategies at once: one backbone serves mechanistically distinct families, and no search loop is required. Beyond zero-shot design, the same framework extends to unseen functional classes or quantitative traits: *fine-tuned in-painting* adapts the model using target-specific SELEX data to produce IGFBP3-binding aptamers that achieve picomolar affinity and rival or surpass SELEX winners, whereas *iterative optimization* uses task-specific predictors to engineer 5′ UTRs that boost protein production of clinically deployed mRNA medicines, with *in vivo* validation in mice showing enhanced anti-spike IgG responses. Together, these applications demonstrate a single framework that generates functional RNAs across known families on demand, while retaining the flexibility to extend to new design objectives by incorporating task-specific data. This establishes multimodal generative learning as a general route from predictive RNA modeling to programmable design of functional RNAs.

## Results

### Learning a multimodal RNA design grammar

We designed RNAverse to learn a shared RNA design space in which sequences, experimentally measured structural profiles, and functional annotations jointly specify and generate RNA molecules (Fig. 1a). The architecture centers on a 30-layer transformer encoder that processes the three input modalities. Sequence and functional labels are tokenized directly, whereas continuous structural scores are discretized into tokens conditioned on assay-specific indicators to account for variations across probing technologies. Resulting modality-specific vectors are fused by element-wise summation before transformer processing, coupling sequence grammar with experimentally grounded structural and functional signals.

**Fig. 1.**
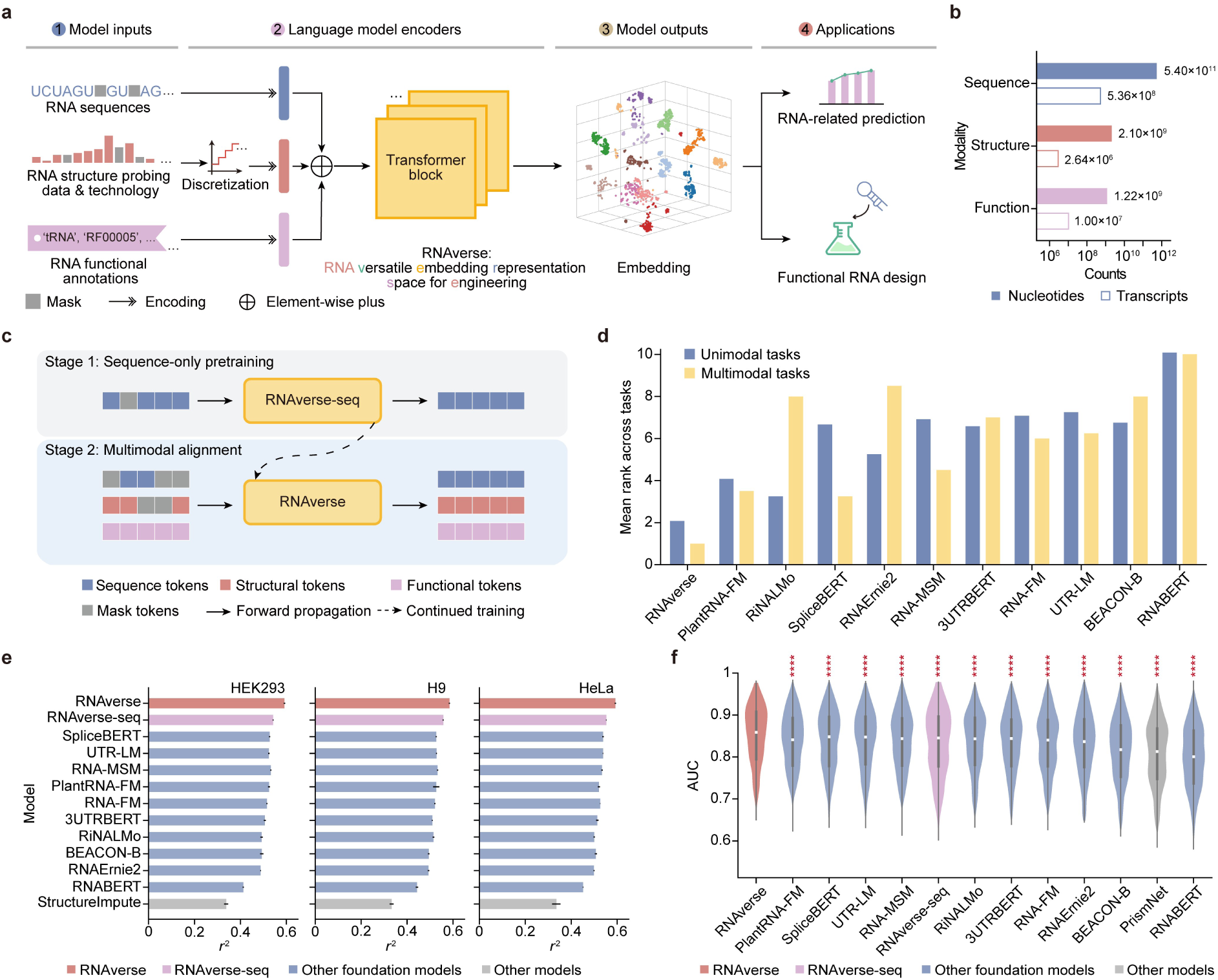
Overview of RNAverse and benchmarking of RNAverse across diverse RNA-related predictive tasks. **a,** Schematic overview of RNAverse, a multimodal RNA foundation model for prediction and functional design. Nucleotide sequences, discretized structure probing data with technology tags, and functional annotations are tokenized and fused into unified embeddings, which are then processed by a 30-layer transformer encoder. The resulting embeddings support downstream prediction and functional RNA design tasks. **b,** Data distribution across RNAverse modalities, showing total nucleotide and transcript counts for the sequence, structure, and function modalities. **c,** Two-stage training strategy. Stage 1 involves masked language modeling on large-scale sequence-only data, followed by Stage 2 multimodal alignment, achieved through the joint reconstruction of masked nucleotide and structural tokens while incorporating functional annotations. **d,** Mean rank of each RNA foundation model across 16 prediction tasks, including 12 unimodal (blue) and 4 multimodal tasks (yellow). **e,** Comparison of SSI performance across three cell lines (HEK293, H9, and HeLa). The performance of RNAverse (pink) and RNAverse-seq (purple) is compared against other foundation models (blue) and the StructureImpute method (gray). Models are ranked by their mean performance on the test set across all six cell lines. Performance is measured as the squared Pearson correlation coefficient (*r*^2^). Validation and test sets were de-duplicated against the Stage 2 alignment corpus. Data are presented as the mean ± s.d from *n* = 3 replicates with different random seeds. **f,** Comparison of RPI performance across RNA-binding proteins (RBPs) (*n* = 172). Violin plots show AUC distributions of RNAverse (pink), RNAverse-seq (purple), other foundation models (blue), and the PrismNet method (gray). Models are ranked by their mean AUC value on the test set across *n* = 172 RBPs. Asterisks indicate statistical significance between RNAverse and other models. *P*-values were determined using a two-sided paired Student’s *t*-test. \**P*≤0.05, \*\**P* ≤ 0.01, \*\*\**P* ≤ 0.001, \*\*\*\**P* ≤ 0.0001; n.s., not significant (*P* > 0.05).

To overcome the intrinsic imbalance between abundant sequence data and scarce structural or functional labels, we curated a large-scale corpus and adopted a two-stage training strategy (Fig. 1b,c). In Stage 1 of sequence-only pretraining, we drew on a non-redundant corpus aggregated from diverse public databases^21–30^ (Extended Data Fig. 1a). This phase captured generalizable evolutionary patterns and compositional sequence features across a broad taxonomic spectrum (Extended Data Fig. 1b). The resulting model, termed RNAverse-seq, served as the backbone for Stage 2 of multimodal alignment, which integrates structure probing data from RASP v2.0^31^ and functional annotations from Rfam^32^ (Extended Data Fig. 1a,c). Here, the model jointly reconstructs masked nucleotides and structural scores conditioned on functional labels, aligning sequence-derived representations with experimental structure and curated function. This fusion improves predictive accuracy and unlocks function-conditioned design ranging from unconstrained *de novo* generation to conditional design guided by arbitrary partial sequence specifications.

Throughout training, RNAverse exhibited robust power-law scaling (Extended Data Fig. 1d). In Stage 1 sequence-only pretraining, scaling was highly consistent (*R*^2^ = 0.996). This trend persisted in Stage 2 multimodal alignment, with *R*^2^ values of 0.979 for sequence reconstruction and 0.965 for structural score prediction. These results show that heterogeneous RNA information can be integrated without sacrificing scalable sequence learning, yielding representations for both generalizable RNA modeling and specification-guided design.

### Multimodal fusion improves RNA prediction most when structure is part of the problem

We benchmarked RNAverse against 10 recent RNA foundation models^33–42^ across 16 prediction tasks: 12 unimodal sequence-based tasks from the BEACON suite^36^ and 4 multimodal tasks combining sequence and structural probing inputs: structural score imputation (SSI)^43^, RNA–protein interaction (RPI) prediction^44^, modification site prediction (MSP), and polyadenylation site (PAS) prediction (Fig. 1d-f, and Extended Data Fig.1e-h). For models available in multiple configurations, we evaluated all variants and reported the best performance (Extended Data Fig.1h; complete results for all 18 model variants are reported in Supplementary Table 4). Briefly, RNAverse achieved the highest mean rank across both unimodal and multimodal benchmarks (Fig. 1d). Strikingly, despite a compact 150-million-parameter architecture, RNAverse outperformed substantially larger models, including the 4.3×-larger RiNALMo^42^ (Fig. 1d).

Performance gains were most pronounced for multimodal tasks demanding a unified view of sequence and structure. We included RNAverse-seq (the Stage 1 sequence-only baseline) as an ablation control. Across all four multimodal benchmarks, RNAverse consistently surpassed RNAverse-seq and all other foundation models, demonstrating that integrating structural information during Stage 2 improves joint sequence–structure reasoning (Fig. 1e,f and Extended Data Fig. 1e-g).

On the SSI task, which imputes missing structural probing data from full sequence and partial structure, RNAverse achieved the highest Pearson correlation coefficient *(r*) across all six cell lines tested, outperforming unimodal models and the task-specific state-of-the-art method, StructureImpute^43^ (Fig. 1e and Extended Data Fig. 1e). On the RPI task, which predicts RBP binding from joint sequence and structural context, RNAverse achieved overall the best area under the receiver operating characteristic curve (AUC) values across 172 RBPs, outperforming all competitors (Fig. 1f). RNAverse likewise led on the MSP and PAS tasks (Extended Data Fig. 1f,g). Across all 16 prediction tasks, RNAverse’s relative performance (z-score relative to all evaluated foundation models) was uncorrelated with test-set similarity to either the full training corpus or the Stage 2 alignment corpus (*r* = −0.04 and −0.05, respectively; Extended Data Fig. 1i), indicating that its performance advantage is unlikely to be explained by test-set similarity to the training data. Together, these results show that multimodal alignment with experimental structural profiles strengthens predictive RNA modeling, particularly for tasks in which structure informs function.

### RNAverse designs PTC-readthrough snoRNAs outperforming rational engineering through zero-shot in-painting

We next asked whether the integrated representations learned through multimodal alignment could be inverted to generate functional RNAs directly from specifications. As a therapeutically demanding testbed, we targeted the repair of premature termination codons (PTCs), genetic lesions underlying ∼11% of inherited diseases and >20% of pathogenic single-nucleotide substitutions^45,46^. We built on the RESTART platform, which employs H/ACA box snoRNAs to guide site-specific pseudouridylation (Ψ) at PTCs and restore protein expression^47^. In RESTART snoRNAs, two internal loops dictate target specificity while the surrounding scaffold governs stability and RNP assembly^47–49^; we reasoned that RNAverse could redesign this scaffold to enhance therapeutic potency.

To do so, we formulated the design as a constrained sequence-completion problem and deployed RNAverse’s generative capability in a zero-shot framework (Fig. 2a). Concretely, the target-specific guide loops were held fixed along with functional specification while the surrounding scaffold nucleotides were masked. The model then in-painted the masked positions, sampling from the conditional distribution shaped during sequence-only pretraining and multimodal alignment. We first applied this strategy to the gACA19 family, generating 500 candidate scaffolds. All 500 scaffolds exhibited lower sequence similarity to natural counterparts than the rationally engineered RESTART-gACA19 construct^47^ (Fig. 2b), indicating that the model explores scaffold space well beyond the reach of human expert design. We randomly selected 20 designs for validation in HEK293T cells. Every variant significantly exceeded the non-targeting control (gCtrl) for PTC-readthrough, with the top variant achieving 14.2-fold and 65.6-fold enhancements in relative eGFP intensity and eGFP-positive cell fraction, respectively. Critically, a substantial fraction also surpassed the rationally engineered RESTART-gACA19 construct: 40% (8/20) and 50% (10/20) outperformed it on intensity and cell-fraction metrics, the best delivering 1.5-fold and 1.3-fold on the two metrics, respectively (Fig. 2c).

**Fig. 2.**
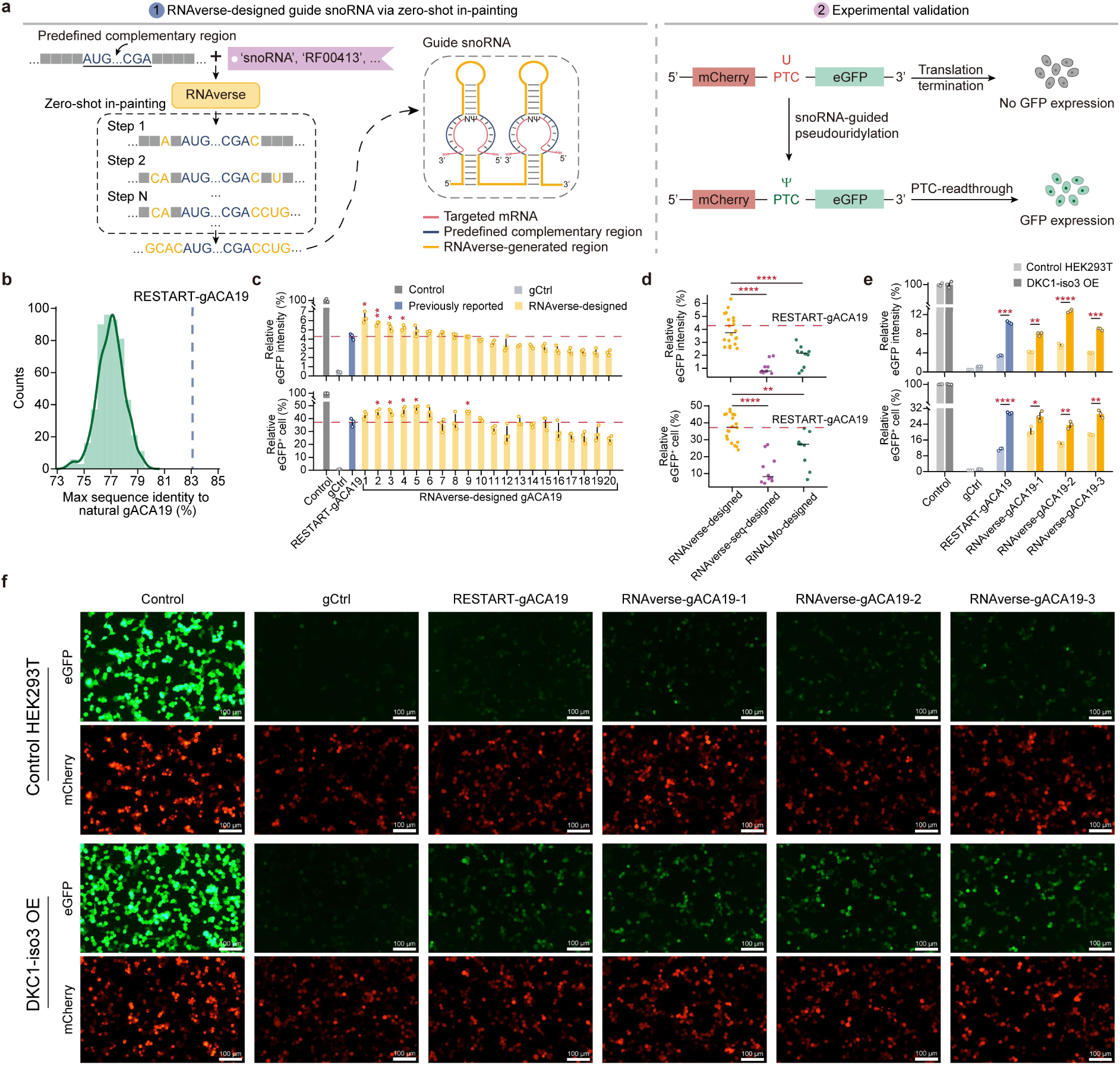
Targeted PTC-readthrough enabled by generatively designed high-potency snoRNA scaffolds. **a,** Schematic of *de novo* PTC-readthrough snoRNA design via RNAverse zero-shot in-painting and subsequent experimental validation. Through a zero-shot in-painting strategy, given functional tokens and partially or fully masked sequences, RNAverse predicts the masked positions based on conditional probabilities, enabling the *de novo* generation of complete functional RNAs. Within the designed guide snoRNA structure, the predefined reverse-complementary internal loops acting as design constraints are shown in dark blue, the RNAverse-generated regions are highlighted in yellow, and the targeted mRNA is depicted in pink. To experimentally validate the generated RNAs, a reporter system is employed where an mRNA contains a premature termination codon (PTC) positioned upstream of an eGFP coding sequence, which normally leads to translation termination and a loss of GFP expression. Upon site-specific pseudouridylation (Ψ) at the PTC guided by the designed snoRNA, translational readthrough is promoted, thereby restoring downstream eGFP expression. **b,** Maximum sequence identity distribution of 500 RNAverse-designed gACA19 variants. For each design, the maximum identity was calculated against all natural gACA19 sequences in the Rfam database. The blue dashed line indicates the maximum sequence identity of the rationally engineered RESTART-gACA19 construct to the same set of natural sequences. **c,** Functional evaluation of PTC-readthrough efficiency for 20 RNAverse-designed gACA19 variants in HEK293T cells. Relative eGFP intensity normalized to the Control group (top) and the relative fraction of eGFP-positive cells (bottom) were measured by flow cytometry. Control (dark grey) represents a reporter with the PTC replaced by the GGU codon to allow constitutive eGFP expression, while gCtrl (light grey) represents a non-targeting guide RNA. RNAverse-designed variants (yellow) are named and ordered by decreasing relative eGFP intensity. Data are presented as the mean ± s.d from *n* = 3 biological replicates. The red dashed line indicates the mean performance of RESTART-gACA19 (blue). Asterisks indicate statistical significance for designs with higher mean values compared to RESTART-gACA19. *P*-values were determined using a two-sided Student’s *t*-test. \**P*≤0.05, \*\**P* ≤ 0.01, \*\*\**P* ≤ 0.001, \*\*\*\**P* ≤ 0.0001; n.s., not significant (*P* > 0.05). **d,** Functional evaluation of PTC-readthrough efficiency for 20 RNAverse-designed (yellow), 10 RNAverse-seq-designed (purple), and 10 RiNALMo-designed (green) gACA19 variants in HEK293T cells. Relative eGFP intensity normalized to the Control group (top) and the relative fraction of eGFP-positive cells (bottom) were measured by flow cytometry. Each point represents the mean of *n* = 3 biological replicates for one design. Black horizontal lines indicate the median of each group. The red dashed line indicates the mean performance of RESTART-gACA19. Asterisks indicate significant differences between RNAverse and each sequence-only baseline (RNAverse-seq and RiNALMo, respectively). *P*-values were determined using a two-sided Student’s *t*-test. \**P*≤0.05, \*\**P* ≤ 0.01, \*\*\**P* ≤ 0.001, \*\*\*\**P* ≤ 0.0001; n.s., not significant (*P* > 0.05). **e,** Effect of DKC1-isoform3 (DKC1-iso3) overexpression on PTC-readthrough efficiency. Relative eGFP intensity (top) and the relative fraction of eGFP-positive cells (bottom) were compared between cells with (dark shading) or without (light shading) DKC1-iso3 overexpression. Data are presented as the mean ± s.d from *n* = 3 biological replicates. *P*-values were determined using a two-sided Student’s *t*-test. \**P*≤0.05, \*\**P* ≤ 0.01, \*\*\**P* ≤ 0.001, \*\*\*\**P* ≤ 0.0001, n.s., not significant (*P* > 0.05). **f,** Representative fluorescence images of the reporter system in HEK293T cells. The top and bottom panels show control cells and cells with DKC1-iso3 overexpression, respectively. eGFP (green) and mCherry (red, internal control) signals are shown for the indicated groups. Scale bars, 100 μm.

To assess the contribution of Stage 2 multimodal alignment to the model’s design capability, we repeated the design procedure with two sequence-only baselines (RNAverse-seq and RiNALMo) as an ablation. Both baselines were fine-tuned on the same gACA19 family sequences using the same masked language modeling objective and then used to generate scaffolds under identical sequence constraints. RNAverse designs were significantly superior to both baselines, with most designs exceeding the best performance achieved by each baseline: 20/20 and 17/20 vs. RNAverse-seq, and 14/20 and 11/20 vs. RiNALMo, for the intensity and cell-fraction metrics, respectively (Fig. 2d and Extended Data Fig. 2a), supporting that multimodal alignment underlies RNAverse’s performance for snoRNA scaffold design.

Fluorescence imaging confirmed robust reporter activation, consistent with efficient PTC readthrough (Extended Data Fig. 2b). We next asked whether the designs could synergize with known system optimizations. We focused on DKC1, the catalytic Ψ synthase in the H/ACA snoRNP complex^50^, whose minor isoform DKC1-isoform3 (DKC1-iso3) reportedly boosts RESTART-mediated PTC readthrough^47^. We found that overexpressing DKC1-iso3 further increased readthrough for all three top RNAverse designs (Fig. 2e,f), supporting that the generated scaffolds engage the native modification machinery.

To assess generality across additional families, we extended the approach to gACA2b and gACA36. As with gACA19, RNAverse-generated sequences diverged substantially from natural counterparts (Extended Data Fig. 2c,d). For gACA2b, several RNAverse designs significantly outperformed the rational construct, with the top candidate achieving a 1.4-fold increase in readthrough efficiency by relative eGFP intensity (Extended Data Fig. 2e). For gACA36, the RNAverse designs matched the human-optimized counterpart, and every tested variant (20/20) exhibited significant activity relative to the non-targeting control (Extended Data Fig. 2f). Together, these results establish RNAverse’s zero-shot, constraint-aware in-painting as a generalizable engine for therapeutic RNA editing, requiring only a small number of experimental validations to identify variants that substantially outperform existing methods.

### RNAverse composes modular RNA activities into chimeric aptazymes for ligand-controlled gene regulation

Having shown that RNAverse can design within a single functional class, we next tested whether it could take a further step: to compose multiple coupled biophysical activities into a single synthetic RNA device. As a test of this modular-composition capability, we targeted RNA aptazymes, non-natural chimeric fusions of a ligand-binding aptamer with a self-cleaving ribozyme^51,52^ used for ligand-controlled gene regulation for precision gene therapy^53^ and metabolic engineering^54^. Here, ligand recognition, ribozyme catalysis, and allosteric communication must be specified simultaneously rather than optimized as isolated functions. Lacking natural templates, state-of-the-art aptazymes have historically required large-scale screening of randomized functional motifs of restricted length^7–9^.

To test whether RNAverse could bypass this screening bottleneck and generate functional aptazyme chimeras, we embedded a designed guanine aptamer within a self-cleaving hammerhead ribozyme (HHR) context such that ligand binding stabilizes local structure and sterically disrupts a required tertiary contact, thereby inhibiting self-cleavage^55–57^. As with the snoRNA design, we used zero-shot in-painting under a composite specification: along with the functional annotation, we fixed an anchor sequence within the ribozyme loop to serve as the stem for the guanine aptamer and used RNAverse to in-paint the remaining scaffold (including the communication module), yielding complete guanine-responsive chimeras without task-specific fine-tuning (Fig. 3a). All 500 generated candidates exhibited lower sequence similarity to natural guanine riboswitches in Rfam than the riboswitch portion of GuaB4HHR, the top variant from a prior HTS^9^ based on the same architecture (Fig. 3b), indicating that RNAverse was not simply retrieving natural riboswitch-like solutions.

**Fig. 3.**
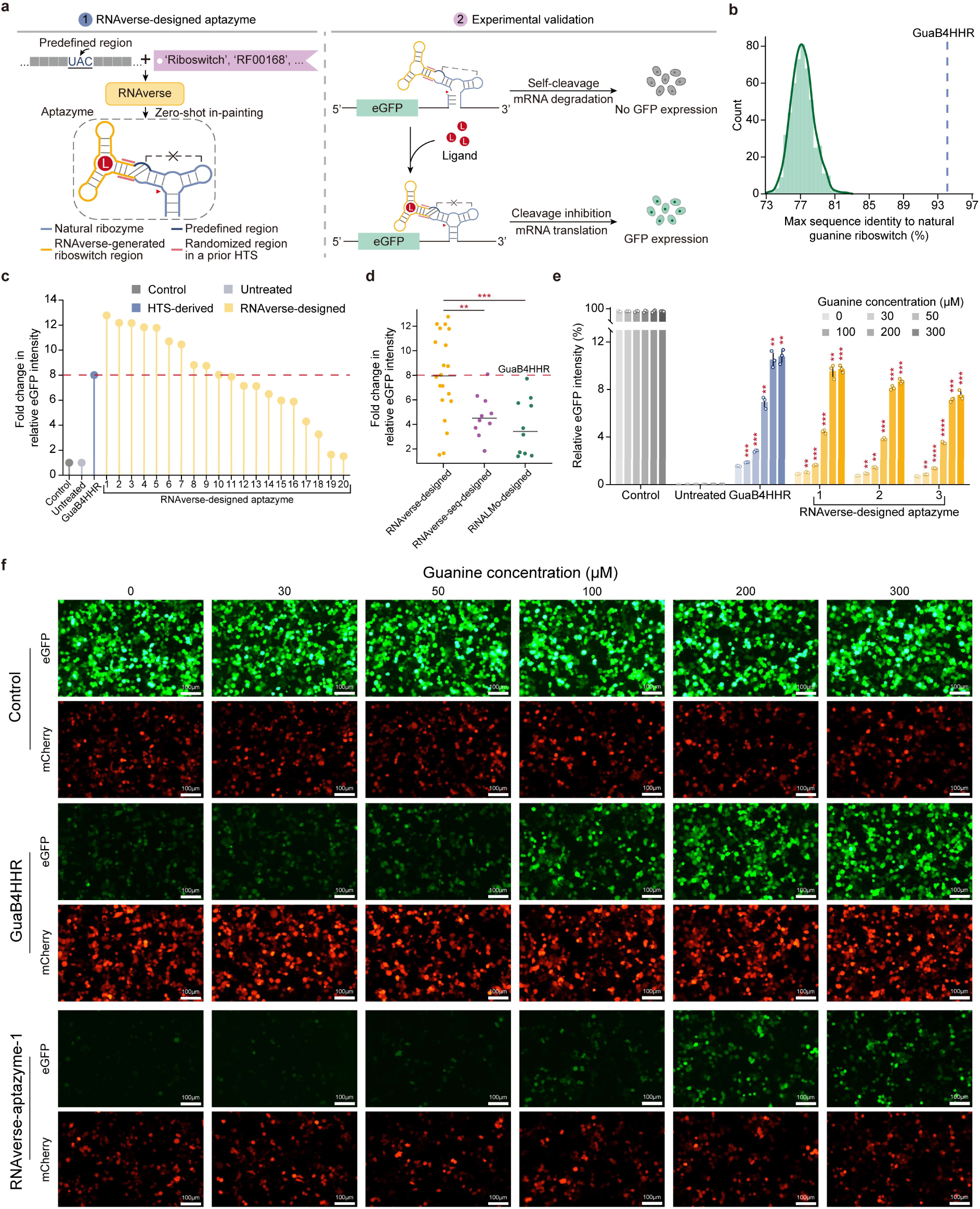
Generative design of high-performance guanine-responsive aptazymes. **a,** Schematic of *de novo* ligand-responsive aptazyme design via RNAverse zero-shot in-painting and subsequent experimental validation. The aptazyme, which is composed of a hammerhead ribozyme fused to a guanine aptamer, is generated *de novo* by RNAverse utilizing the zero-shot in-painting strategy. Within the aptazyme diagram, the hammerhead ribozyme scaffold is shown in light blue, while predefined anchor sequences within the ribozyme loop acting as design constraints in the stem of guanine aptamer are colored in dark blue. RNAverse-generated regions are highlighted in yellow, and pink lines indicate the regions randomized in a prior high-throughput screening (HTS) library9 for performance benchmarking. To experimentally validate the designed aptazymes, a ligand-responsive reporter system is employed where the aptazyme is positioned in the 3′ UTR of an eGFP coding sequence. In the absence of guanine, ribozyme self-cleavage (red triangle) promotes mRNA degradation and suppresses eGFP expression. Upon guanine binding, the ribozyme tertiary loop-loop interactions (black dashed lines) are disrupted, which inhibits cleavage and restores downstream eGFP expression. **b,** Maximum sequence identity distribution of 500 RNAverse-designed guanine riboswitch variants. For each design, the maximum identity was calculated against all natural guanine riboswitch sequences in the Rfam database. The blue dashed line indicates the maximum sequence identity of the riboswitch portion of the HTS-derived top variant GuaB4HHR to the same set of natural sequences. **c,** Fold change of eGFP expression for 20 RNAverse-designed aptazymes, calculated as the ratio of mean relative eGFP intensity at 300 μM versus 0 μM guanine from *n* = 3 biological replicates. RNAverse-designed variants are named and ordered by decreasing fold change. Control denotes a non-aptazyme reporter, Untreated indicates untransfected cells, and GuaB4HHR is a top-performing HTS-derived variant (the red dashed line). **d,** Fold change of eGFP expression for 20 RNAverse-designed (yellow), 10 RNAverse-seq-designed (purple), and 10 RiNALMo-designed (green) aptazymes, calculated as the ratio of mean relative eGFP intensity at 300 μM versus 0 μM guanine from n = 3 biological replicates. Black horizontal lines indicate the median of each group. The red dashed line indicates the performance of GuaB4HHR. Asterisks indicate significant differences between RNAverse and each sequence-only baseline (RNAverse-seq and RiNALMo, respectively). *P*-values were determined using a two-sided Student’s *t*-test. \**P*≤0.05, \*\**P* ≤ 0.01, \*\*\**P* ≤ 0.001, \*\*\*\**P* ≤ 0.0001; n.s., not significant (*P* > 0.05). **e,** Dose-response evaluation of top-performing RNAverse-designed aptazymes. Relative eGFP intensity for GuaB4HHR and the top three RNAverse-designed variants was measured across increasing guanine concentrations (0, 30, 50, 100, 200, and 300 μM). For each variant, the bars represent ligand concentrations from low to high (from light shading to dark shading). Data are presented as the mean ± s.d from *n* = 3 biological replicates. Asterisks indicate statistical significance for each concentration versus the 0 μM group within each variant. *P*-values were determined using a two-sided Student’s *t*-test. \**P*≤0.05, \*\**P* ≤ 0.01, \*\*\**P* ≤ 0.001, \*\*\*\**P* ≤ 0.0001; n.s., not significant (*P* > 0.05). **f,** Representative fluorescence images of aptazyme-regulated eGFP expression in HEK293T cells. eGFP (green) and mCherry (red, internal control) signals are shown in cells treated with increasing concentrations of guanine for GuaB4HHR (top) and RNAverse-aptazyme-1 (bottom). Scale bar, 100 μm.

We randomly selected 20 RNAverse designs for testing in HEK293T cells and benchmarked them against GuaB4HHR. All 20 exhibited significant ligand-dependent eGFP induction (Extended Data Fig. 3a), demonstrating that RNAverse can directly compose functional aptazyme chimeras from coupled RNA activities without large-scale screening. Strikingly, 50% (10/20) of RNAverse chimeras achieved higher fold eGFP induction than GuaB4HHR. The best RNAverse design reached a 12.8-fold induction upon guanine treatment, a 1.6-fold improvement over GuaB4HHR, driven primarily by tighter off-state expression with comparable on-state output (Fig. 3c and Extended Data Fig. 3a).

As in the snoRNA task, we benchmarked against the two sequence-only baselines to isolate the contribution of multimodal alignment. RNAverse designs showed significantly higher fold change than both baselines, with a substantial fraction exceeding the best performance of each baseline: 9/20 vs. RNAverse-seq, and 11/20 vs. RiNALMo, respectively (Fig. 3d and Extended Data Fig. 3a). Together with the snoRNA results, these findings support a reproducible contribution of multimodal alignment to zero-shot RNA device design across two distinct functional settings.

Dose-response measurements for the top three RNAverse designs confirmed sensitive, concentration-dependent regulation across 0–300 μM guanine by flow cytometry and fluorescence microscopy (Fig. 3e,f and Extended Data Fig. 3b). Together, these results establish RNAverse’s zero-shot, constraint-aware in-painting as a route to *de novo* chimeric RNA devices that satisfy coupled biophysical constraints simultaneously, reaching functional regimes beyond randomized screening with substantially reduced experimental validation.

### RNAverse design extends to unseen functional classes of high-affinity aptamers via fine-tuned in-painting

The preceding designs redeployed functional classes already represented during multimodal alignment, for which RNAverse had already learned the relevant priors. We next asked whether this design grammar could extend to an entirely unseen functional class, where such priors must instead be acquired *de novo* from task-specific data. As a test case, we targeted IGFBP3-binding aptamers, a class absent from the alignment corpus, whose binding determinants are encoded only in specialized experimental measurements rather than in public functional databases. To acquire task-specific priors, we fine-tuned RNAverse on 13,802 sequences from the final round of a SELEX experiment^58^ using a masked language modeling objective, introducing a dedicated **`[IGFBP3_aptamer]`** token to specify the task. The resulting RNAverse-aptamer was used to generate 36-nt aptamers through iterative in-painting (Fig. 4a). We term this strategy “fine-tuned in-painting”: it extends RNAverse’s in-painting to functional classes outside the alignment corpus by adapting pretrained priors through task-specific updates.

**Fig. 4.**
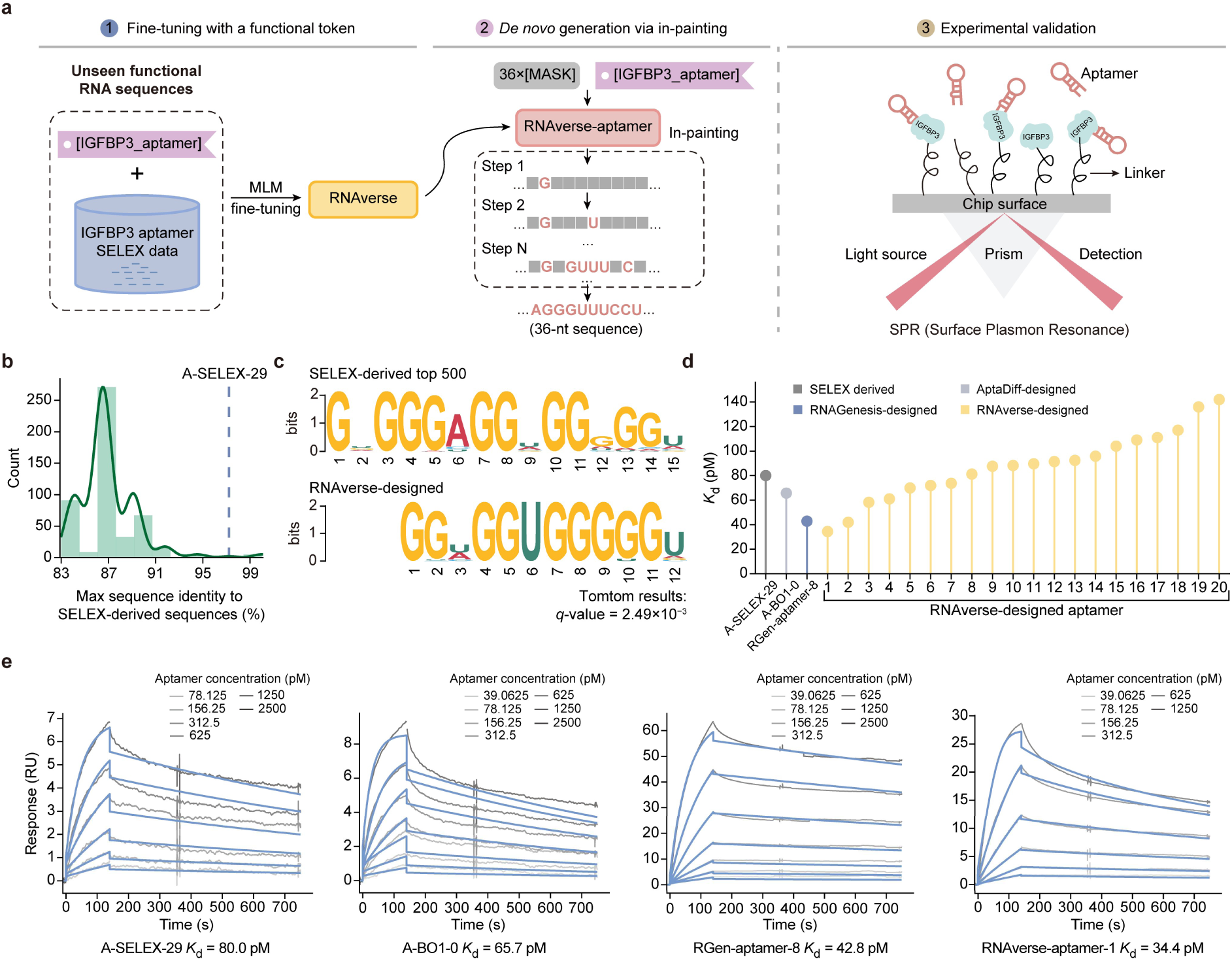
*De novo* design of high-affinity IGFBP3 aptamers through fine-tuned in-painting. **a,** Schematic of IGFBP3 aptamer design via RNAverse fine-tuned in-painting and subsequent experimental validation. Specifically, RNAverse was first fine-tuned on SELEX-derived sequences via a masked language modeling (MLM) objective using a novel functional token `[IGFBP3_aptamer]` to obtain RNAverse-aptamer. This specialized model was utilized for the generation of 36-nt aptamer regions via in-painting, followed by experimental validation of binding affinity using surface plasmon resonance (SPR). **b,** Maximum sequence identity distribution of 500 RNAverse-designed IGFBP3 aptamers. For each design, the maximum identity was calculated against all 13,802 final-round SELEX sequences. The blue dashed line indicates the maximum sequence identity between the top SELEX-derived A-SELEX-29 and its closest neighbor among the other final-round SELEX sequences. **c,** Comparison of enriched sequence motifs between the top 500 SELEX-derived sequences and 500 RNAverse-designed sequences. Motif alignment and similarity were assessed using Tomtom, with *q*-values indicating statistical significance. **d,** Binding affinities (*K*_d_) of RNAverse-designed aptamers compared to benchmark sequences. Bar plots show the *K*_d_ (pM) determined by SPR. Benchmark sequences include the top variant from SELEX-derived (A-SELEX-29, dark gray), AptaDiff-designed (A-BO1-0, light gray), and RNAGenesis-designed (RGen-aptamer-8, blue). RNAverse-designed variants (yellow) are named and ordered by increasing *K*_d_. **e,** SPR sensorgrams for A-SELEX-29, A-BO1-0, RGen-aptamer-8, and the top RNAverse-designed aptamer (RNAverse-aptamer-1). Curves in varying shades of gray represent experimental binding responses (RU) at increasing analyte concentrations, and blue curves indicate the kinetic fits used for *K_d_* calculation.

The median maximum identity of the 500 RNAverse-designed aptamers to any final-round SELEX sequence was 86.5%, lower than the 97.2% identity between A-SELEX-29 (the top SELEX-derived aptamer) and its closest final-round SELEX neighbor (Fig. 4b). This indicates that fine-tuning had not simply memorized or retrieved the SELEX pool. At the same time, motif-enrichment analysis revealed strong overlap with motifs from top-500 performing SELEX sequences (Tomtom^59^ *q*-value = 2.49 × 10^-3^), suggesting that the model had internalized the binding grammar of the class (Fig. 4c).

We then randomly selected 20 RNAverse-designed aptamers for surface plasmon resonance (SPR) validation. None of these 20 were identical to any sequence in the final-round SELEX library (Raw data of Fig. 4b). All 20 bound IGFBP3 with picomolar affinity (*K*_d_ = 34.4–142 pM; Fig. 4d,e and Extended Data Fig. 4a). Strikingly, the best RNAverse design (*K*_d_ = 34.4 pM) was the highest-affinity binder across all tested sequences, surpassing the top aptamers from SELEX (A-SELEX-29, 80.0 pM), AptaDiff^58^ (A-BO1-0, 65.7 pM), and RNAGenesis (RGen-aptamer-8, 42.8 pM). When benchmarked against top-performing sequences from previous methods, 35% (7/20) of designs outperformed the best SELEX-derived aptamer, while 20% (4/20) and 10% (2/20) surpassed the top designs from AptaDiff and RNAGenesis, respectively (Fig. 4d,e and Extended Data Fig. 4a). Thus, even for functional classes outside its alignment corpus, RNAverse converts pretrained priors into high-potency RNAs through lightweight fine-tuning, expanding its design reach.

### From categorical to quantitative RNA design: RNAverse optimizes 5′ UTRs to boost therapeutic protein expression

The second extension moves from categorical classes to quantitative traits. Beyond categorical RNA classes, many therapeutic applications require quantitative tuning of continuous functional properties. Here the challenge is not reaching an unseen functional class but optimizing a continuous phenotype: an objective for which no categorical prior exists. As a clinically motivated testbed, we used RNAverse to engineer 5′ UTRs for translational control by coupling high-throughput quantitative phenotyping with predictor-guided sequence search. We constructed a library of constructs containing 5′ UTR fragments from human and viral sources^60^, experimentally profiled their structures by icSHAPE-MaP^61^, and quantified their translation by high-throughput reporter-based measurements in HEK293T cells (Fig. 5a). After quality filtering, this newly generated dataset comprises high-quality structural and translational data for over 17,000 fragments (Extended Data Fig. 5a-c), with strong concordance to established structural models and replicate libraries (Extended Data Fig. 5d-f). Pooled protein estimates also agreed closely with single-construct fluorescence measurements (Fig. 5b). The profiling recapitulated known regulatory logic: Kozak-enriched fragments boosted protein production^62,63^, whereas RNA G-quadruplex (RG4)-containing fragments suppressed it^64,65^ (Extended Data Fig. 5g), establishing this resource as a biologically faithful basis for modeling structure-dependent translational control by 5′ UTRs.

**Fig. 5.**
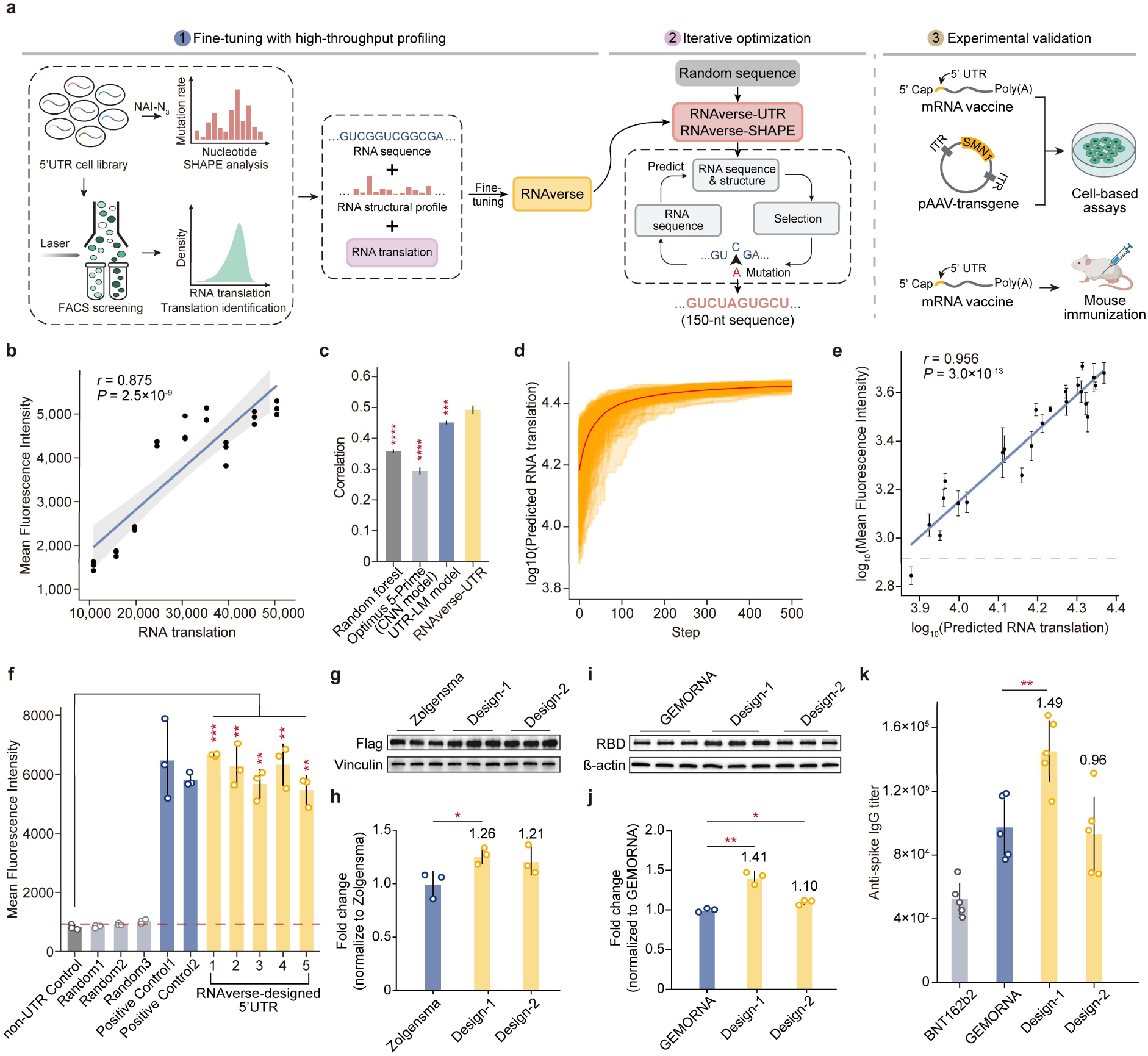
RNAverse-enabled translational optimization leveraging high-throughput 5′ UTR structure and translation profiling. **a,** Schematic of high-translation 5′ UTR design via RNAverse iterative optimization. A 150-nt 5′ UTR library was profiled by high-throughput SHAPE probing to quantify RNA secondary structure, alongside high-throughput eGFP reporter–based measurements of RNA translation. Leveraging this dataset, RNAverse was fine-tuned to obtain RNAverse-UTR, which integrates sequence and structural features of the 150-nt 5′ UTR and the initial 50-nt of CDS to predict RNA translation. This model was then embedded within an iterative optimization strategy, where it evaluated translational performance based on sequences and their predicted SHAPE profiles to guide successive rounds of mutation and selection, enabling the design of high-translation 5′ UTRs. We validated this strategy on two clinically relevant mRNA contexts: the 5′ UTR design for the BNT162b2 COVID-19 vaccine and an AAV9-delivered gene therapy (Zolgensma). For the BNT162b2 context, *in vivo* validation was performed by immunizing mice with mRNA bearing designed 5′ UTRs. Mouse and syringe illustrations were created with BioRender. **b,** Relationship between individual reporter measurements (mean eGFP fluorescence intensity, MFI) and pooled profiling estimates (RNA translation) for selected 5′ UTR variants. Pearson correlation coefficient (*r*) and the *P* value are shown. **c,** Predictive performance of RNAverse-UTR and other models. Correlation coefficients between predicted and measured values are shown for RNA translation. Data are presented as the mean ± s.d over *n* = 5 cross-validation folds. Asterisks indicate statistical significance between RNAverse-UTR and other models. *P*-values were determined using a two-sided paired Student’s *t*-test. \**P*≤ 0.05, \*\**P* ≤ 0.01, \*\*\**P* ≤ 0.001, \*\*\*\**P* ≤ 0.0001; n.s., not significant (*P* > 0.05). **d,** Predicted RNA translation of generated 5′ UTRs across successive iterations of the genetic algorithm. **e,** Scatter plot showing the correlation between predicted RNA translation versus measured MFI in reporter assays for RNAverse-designed 5′ UTRs. Pearson correlation coefficient (*r*) and the *P* value are shown. **f,** MFI of constructs containing RNAverse-designed 5′ UTRs, alongside non-5′ UTR control, randomly generated 5′ UTRs (Random 1–3), and previously identified positive controls (positive control 1 from our initial screen, positive control 2 from previously reported). Data are presented as the mean ± s.d from *n* = 3 biological replicates. The blue dashed line indicates the mean performance of the non-5′ UTR control. Asterisks indicate statistical significance between RNAverse-designed 5′ UTRs and the non-UTR control. *P*-values were determined using a two-sided Student’s *t*-test. \**P*≤0.05, \*\**P* ≤ 0.01, \*\*\**P* ≤ 0.001, \*\*\*\**P* ≤ 0.0001; n.s., not significant (*P* > 0.05). **g,** Representative western blot showing protein expression from Zolgensma constructs bearing original 5′ UTR and RNAverse-UTR-designed 5′ UTRs (Design-1 and Design-2). Vinculin was used as a loading control. **h,** Densitometric quantification of western blot signals. Band intensities were normalized to vinculin and then to the original Zolgensma 5′ UTR reference. Data are presented as mean ± s.d. from *n* = 3 biological replicates. *P*-values were determined using a two-sided Student’s *t*-test. \**P* ≤ 0.05, \*\**P* ≤ 0.01, \*\*\**P* ≤ 0.001, \*\*\*\**P* ≤ 0.0001; n.s., not significant (*P* > 0.05). **i,** Representative western blot showing spike protein expression from BNT162b2 mRNA constructs bearing 5′ UTRs designed by GEMORNA or RNAverse-UTR (Design-1 and Design-2). β-actin was used as a loading control. **j,** Densitometric quantification of western blot signals. Band intensities were normalized to β-actin and then to the GEMORNA-designed BNT162b2 5′ UTR reference. Data are presented as mean ± s.d. from *n* = 3 biological replicates. *P*-values were determined using a two-sided Student’s *t*-test. \**P*≤ 0.05, \*\**P* ≤ 0.01, \*\*\**P* ≤ 0.001, \*\*\*\**P* ≤ 0.0001; n.s., not significant (*P* > 0.05). **k,** Comparison of anti-spike immunoglobulin G (IgG) titer on day 21 at a 10 μg dose. Data are presented as mean ± s.d. from *n* = 5 mice per group. *P*-values were determined using a two-sided Student’s *t*-test. \**P*≤0.05, \*\**P* ≤ 0.01, \*\*\**P* ≤ 0.001, \*\*\*\**P* ≤ 0.0001; n.s., not significant (*P* > 0.05).

Beyond well-defined motifs, 5′ UTRs are governed by distributed sequence features and dynamic structural constraints^66,67^, so effective optimization requires a predictor that captures this regulatory complexity rather than a catalog of individual elements. To build one, we coupled RNAverse’s predictive modeling with a genetic algorithm (Fig. 5a): the backbone is fine-tuned into task-specific predictors, which then guide sequence search. We first fine-tuned RNAverse on our profiling dataset to create RNAverse-UTR, which predicts translational output from joint sequence-structure features. RNAverse-UTR outperformed existing predictors, including UTR-LM^35^, Optimus 5-Prime^68^, and a random forest model^69^, all re-trained on the same profiling dataset using parameters reported in their original studies (Fig. 5c). Ablation experiments showed that structural information contributed a modest but reproducible improvement on this 5′ UTR prediction task (Extended Data Fig. 5h). To generalize structural context and enable cell-type-aware design, we also fine-tuned RNAverse to predict RNA structural profiles directly from sequence, yielding RNAverse-SHAPE, which surpassed StructureImpute^43^ in recapitulating experimental probing data (Extended Data Fig. 5i,j).

Leveraging these two predictors, RNAverse-UTR and RNAverse-SHAPE, the iterative optimization strategy searched the vast 5′ UTR space for maximal protein expression. We observed that predicted GFP output improved monotonically over successive iterations (Fig. 5d), and experimental tests showed a strong correlation between predicted protein production and fluorescence measurements for designed sequences (Fig. 5e). Top RNAverse-designed 5′ UTRs outperformed randomly generated sequences and slightly exceeded two high-expression controls, one previously reported^69^ and one from our initial screen, indicating that this strategy navigates vast sequence spaces beyond the physical limits of experimental libraries. (Fig. 5f).

We then put this workflow to the test on two clinically deployed mRNAs whose efficacy depends directly on per-transcript protein output: Zolgensma, an AAV9-delivered gene therapy for spinal muscular atrophy (SMA), and the BNT162b2 COVID-19 mRNA vaccine. For Zolgensma, where higher SMN output could broaden the therapeutic window of this single-dose treatment, the top design improved protein production by ∼26% over the original 5′ UTR (Fig. 5g,h). For BNT162b2, the clinical 5′ UTR is derived from human α-globin^70^; even modest gains in expression efficiency translate into lower effective doses or broader access at fixed manufacturing capacity. We generated 5′ UTR variants while holding the coding sequence and length fixed, then validated top candidates by reporter assay. The best RNAverse designs achieved ∼41% higher protein production than the GEMORNA-designed 5′ UTR reference^15^ (Fig. 5i,j), which itself produces 93% more protein than the original BNT162b2 (Extended Data Fig. 5k); the top design therefore yields ∼2.7-fold higher protein output than the original clinical construct (computed from the two measured ratios, 1.93 × 1.41, in this reporter assay). To assess *in vivo* humoral responses, mice were immunized with BNT162b2 spike mRNA bearing the original, GEMORNA-designed, or RNAverse-designed 5′ UTRs. At day 21, RNAverse Design1 elicited significantly higher anti-spike IgG titers than both BNT162b2 (2.8-fold) and GEMORNA (1.5-fold) (*n* = 5 mice per group, 10 μg, ALC-0315-based LNP, i.m.), while Design2 showed a 1.8 -fold increase over BNT162b2 (Fig. 5k and Extended Data Fig. 5l). Together, these results extend RNAverse from categorical generation to quantitative optimization, enabling precise translational tuning of therapeutically relevant mRNAs.

## Discussion

These results reframe RNA engineering as a problem of specification. By learning the coupled sequence-structure-function landscape across modalities, RNAverse designs functional RNAs from mechanistically distinct classes directly from stated intent, rather than discovering them by screening. By building a shared representation that connects RNA families and integrates complementary modality-specific information, RNAverse is not merely a more accurate RNA model but a programmable route from desired function to experimentally validated molecules. We show this across three design regimes. Using zero-shot in-painting, RNAverse generated PTC-readthrough snoRNAs and ligand-responsive aptazymes directly from the pretrained backbone. These are only two instances of a much larger design space: thousands of annotated RNA families and millions of modular device combinations are, in principle, accessible to the same backbone without task-specific training. How far the strategy generalizes and at what hit rates remain to be mapped. In the second and third regimes, where tasks demand information beyond the pretrained priors, RNAverse required only lightweight adaptation: fine-tuning on a small target-based prior to generate high-affinity aptamers for an unseen class, and task-specific predictors to guide iterative optimization of translation-tuning 5’ UTRs. In every case fewer than two dozen designs were sufficient to match — and in most cases surpass — the best constructs from rational engineering or large-scale screening. That this consistency holds across distinct mechanisms provides strong evidence that RNA engineering can move from searching libraries for functional molecules to declaring molecular intent and realizing it by design.

Large-scale efforts such as Ribonanza have leveraged crowdsourced structure probing data to advance predictive RNA modeling^71^. RNAverse builds on this progress by multimodal fusion, and in particular, on treating functional annotations as explicit conditioning signals rather than mere evaluation labels: a choice decisive for cross-family generalization. Combined with structural probing data, these annotations let the model learn the semantics of functional families directly, sharing parameters across classes and specifying design intent at inference. Ablation experiments with sequence-only baseline models confirmed that Stage 2 multimodal alignment substantially improves zero-shot design capability, with RNAverse designs consistently outperforming baselines fine-tuned on the same corresponding family sequences. Similar to recent multimodal generative frameworks in protein engineering, such as ESM3^72^, RNAverse suggests that integrating complementary modality-specific information into a shared representation can enable specification-guided biomolecular design. Within one a single framework, RNAverse supports three complementary regimes spanning the full design spectrum. These range from redeploying priors learned during multimodal alignment (zero-shot in-painting), to extending to functional classes never seen in training (fine-tuned in-painting), to tuning continuous properties that lack discrete labels (iterative optimization). Whereas prior generative approaches have operated one family or task at a time, these regimes coexist within one model. Notably, unlike generative methods restricted to global latent-space sampling, RNAverse can infill defined functional segments (*e.g.*, guide loops in snoRNAs, chimeric junctions in aptazymes) within fixed scaffolds, enabling the segment-level control.

We note that in prediction tasks, incorporating structural information improved accuracy over sequence-only models, though these gains were task-dependent: structural information proved most informative when structural state is directly measured and mechanistically linked to function. This pattern shaped design as much as prediction: the major generative design tasks did not uniformly require explicit structural inputs. For well-structured non-coding RNAs such as snoRNAs and riboswitches, evolutionary constraints and family-level annotations can encode substantial information about functional determinants^32,73^. For regulatory elements such as 5′ UTRs, whose functional folds are often transient and poorly captured by consensus structures^66,67^, experimental structure probing data provides an improvement as an additional signal. These results suggest that multimodal learning is most useful when complementary modalities are matched to the biological information required by the task, rather than assuming that every design problem benefits equally from every modality.

Several limitations remain. First, although our two-stage training strategy mitigates data scarcity, structural and functional modalities remain disproportionately sparse relative to the vast repertoire of unlabeled genomic sequences. This sparsity likely explains why structural information provides only a modest predictive gain in the current 5′ UTR optimization pipeline, despite its known importance in translational regulation. Expanding high-quality and condition-resolved structure/function atlases would strengthen RNAverse’s ability to capture subtle conformational dynamics and to improve its accuracy across cellular environments. Second, the current functional annotations, primarily derived from Rfam, provide a high-level categorical framework but lack the granularity required for more nuanced tasks. Incorporating higher-resolution functional layers, such as RNA-protein interactomes, site-specific epitranscriptomic modifications (e.g., m^6^A), and *in vivo* kinetic rates, will be essential for advancing toward more precise, compositionally complex generative tasks. As richer functional and structural atlases become available, we expect the need for task-specific fine-tuning to decrease, with more design tasks becoming accessible in a zero-shot or few-shot manner.

In sum, RNAverse takes a substantive step toward the long-standing challenge of accurate, programmable RNA design, with demonstrated capabilities spanning the major classes of RNA therapeutics: from genetic disease and gene therapy to aptamer drugs and vaccines. As demand grows for programmable RNA engineering, RNAverse and related platforms are poised to accelerate applications in synthetic biology and precision medicine.

## Extended Figure Legends

**Extended Data Fig. 1.**
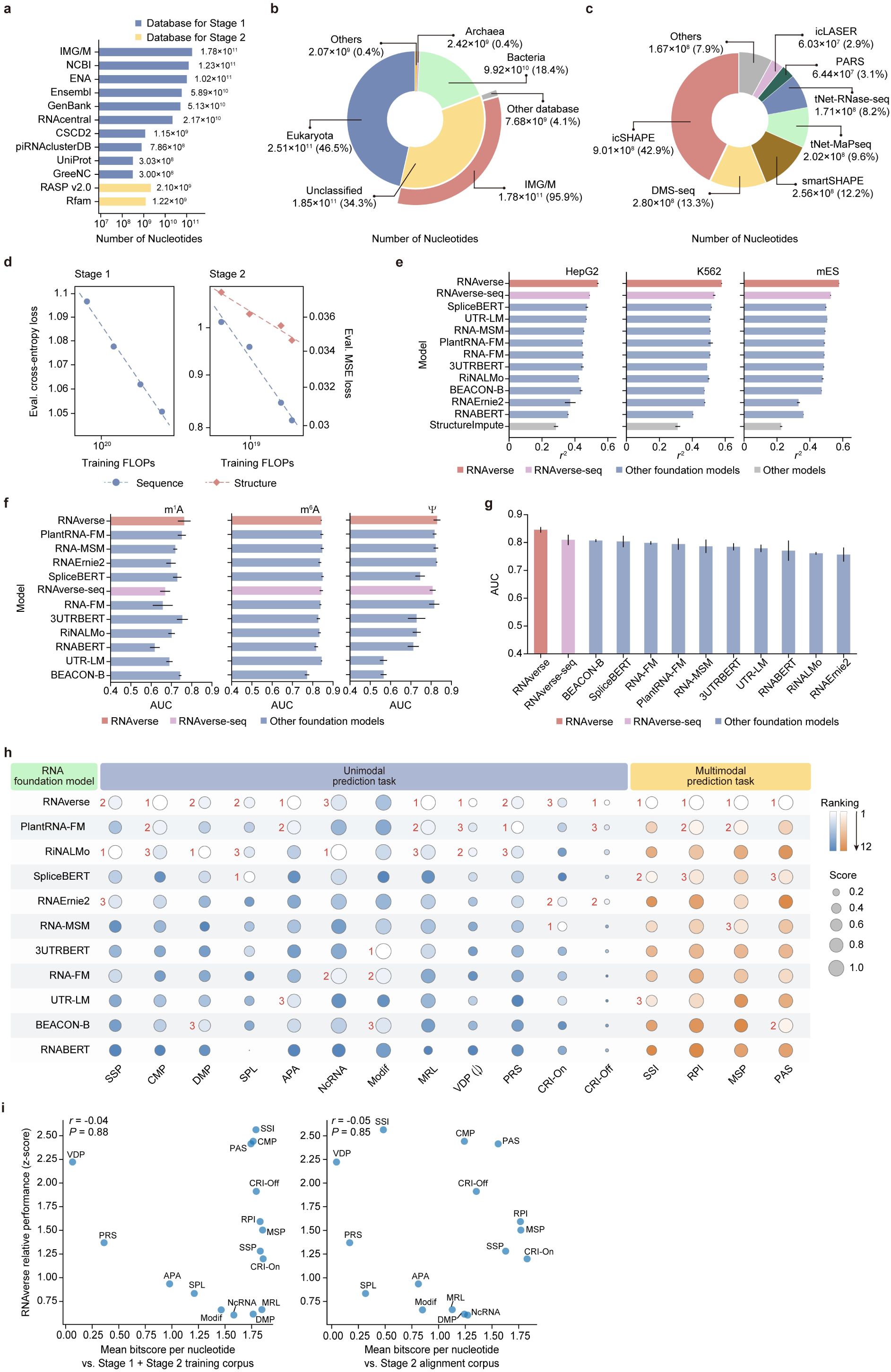
Data composition of the RNAverse training corpus and extended benchmarking. **a,** Data sources for Stage 1 and Stage 2 training. The number of nucleotides across the source database for Stage 1 (sequence-only; blue) and Stage 2 (multimodal; orange) training is shown. **b,** Taxonomic distribution of the training corpus. The pie chart illustrates the proportion of nucleotides assigned to major biological domains, calculated across the Stage 1 and Stage 2 training corpus. The “Unclassified” fraction (34.3%) is predominantly composed of meta-transcriptomic data from the IMG/M database (representing 95.9% of the “Unclassified" fraction). **c,** Experimental probing technology distribution of the Stage 2 alignment corpus. The pie chart illustrates the proportion of nucleotides assigned to different experimental probing technologies. **d,** Scaling laws of RNAverse training. Evaluation loss is plotted as a function of training FLOPs for both stages, employing cross-entropy for sequence reconstruction and mean squared error (MSE) for structural score prediction. **e,** Comparison of SSI performance across three additional cell lines (HepG2, K562, and mES). The performance of RNAverse (pink) and RNAverse-seq (purple) is compared against other foundation models (blue) and the StructureImpute method (gray). Models are ranked by their mean performance on the test set across all six cell lines. Performance is measured as the squared Pearson correlation coefficient (*r*^2^). Validation and test sets were de-duplicated against the Stage 2 alignment corpus. Data are presented as the mean **±** s.d from *n* = 3 replicates with different random seeds. **f,** Comparison of MSP performance across three RNA modification types (m^1^A, m^6^A, and Ψ). The performance of RNAverse (pink) and RNAverse-seq (purple) is compared against other foundation models (blue). Models are ranked by their mean performance on the test set across all three modification types. Data are presented as the mean **±** s.d from *n* = 3 replicates with different random seeds. **g,** Comparison of PAS performance. The performance of RNAverse (pink) and RNAverse-seq (purple) is compared against other foundation models (blue). Models are ranked by their performance on the test set. Data are presented as the mean **±** s.d from *n* = 3 replicates with different random seeds. **h,** Bubble matrix comparing 11 RNA foundation models across 16 prediction tasks, including 14 unimodal (blue) and 4 multimodal tasks (orange). Circle size denotes the performance score, color intensity indicates the relative rank within each task, and red labels mark the top three models within each task. Models are ordered by their mean rank across all tasks. **i,** Relationship between downstream test-set similarity to upstream training corpus and RNAverse’s relative performance across 16 prediction tasks. Each point represents one task. Similarity was measured as mean BLAST bitscore per nucleotide, calculated from the best hit for each test sequence, against (left) the full Stage 1 + Stage 2 corpus and (right) the Stage 2 alignment corpus. RNAverse performance is shown as z-score across all evaluated RNA foundation model variants. Pearson r and *P*-values are shown.

**Extended Data Fig. 2.**
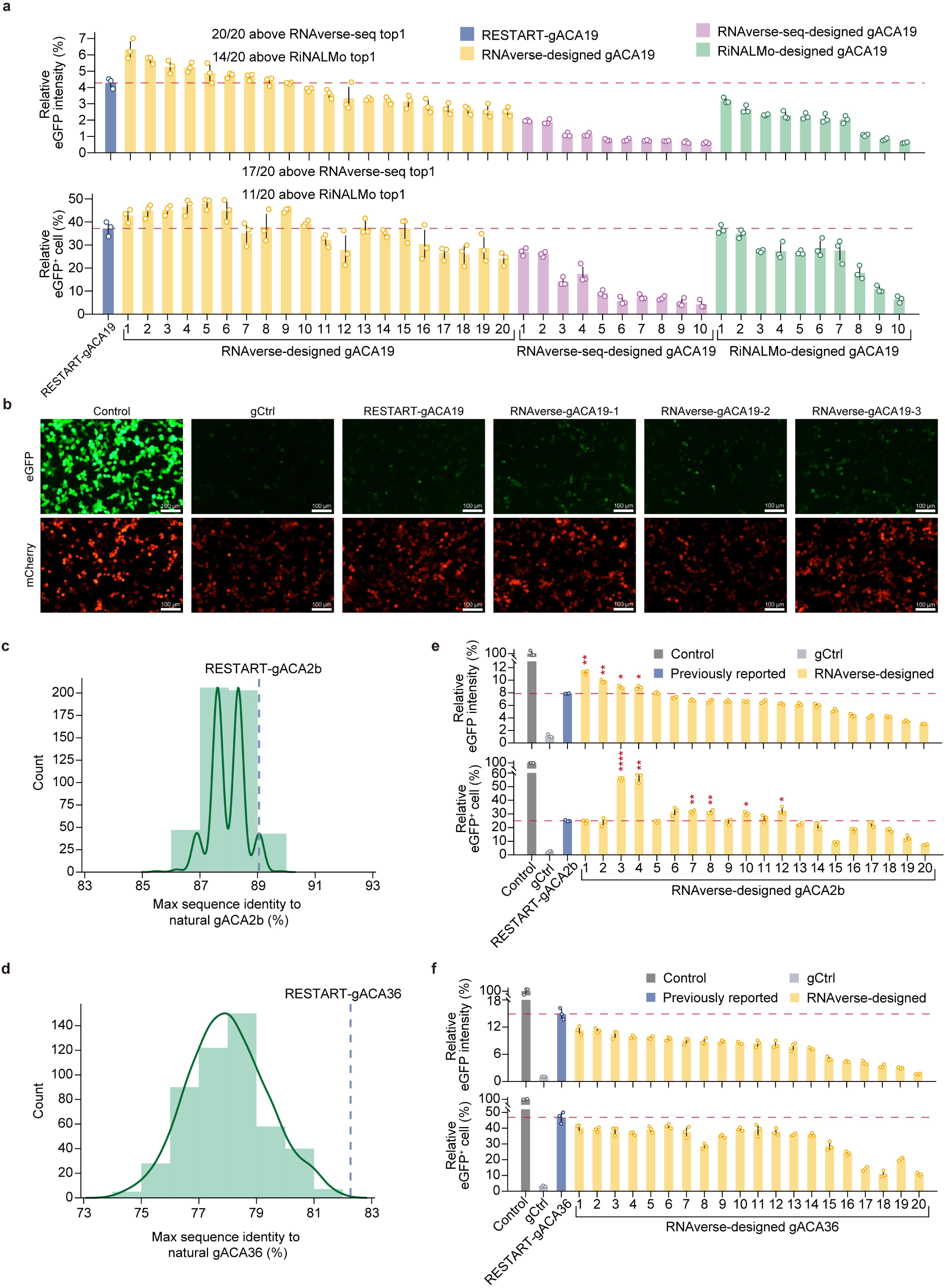
Zero-shot in-painting design framework and extended validation of RNAverse-designed snoRNAs. **a,** Functional evaluation of PTC-readthrough efficiency for 20 RNAverse-designed (yellow), 10 RNAverse-seq-designed (purple), and 10 RiNALMo-designed (green) gACA19 variants in HEK293T cells. Relative eGFP intensity normalized to the Control group (top) and the relative fraction of eGFP-positive cells (bottom) were measured by flow cytometry. Variants within each group are named and ordered by decreasing relative eGFP intensity. Data are presented as the mean ± s.d from *n* = 3 biological replicates. The red dashed line indicates the mean performance of RESTART-gACA19 (blue). RNAverse-designed variants were compared against the top-performing design from each sequence-only baseline: 20/20 and 17/20 designs exceeded the top RNAverse-seq variant, while 14/20 and 11/20 exceeded the top RiNALMo variant in the top and bottom panels, respectively. Bootstrap resampling (10 RNAverse designs sampled without replacement from the 20, 1,000 iterations) assessed robustness to unequal sample sizes; 100% of iterations were significant for all comparisons except RNAverse vs. RiNALMo in the cell-fraction metric, where 99.6% remained significant. Median numbers of RNAverse designs exceeding the baseline top in equal-sample bootstrap: intensity, 10/10 vs. RNAverse-seq and 7/10 vs. RiNALMo; cell fraction, 9/10 vs. RNAverse-seq and 5/10 vs. RiNALMo. **b,** Representative fluorescence images of the reporter system in HEK293T cells. eGFP (green) and mCherry (red, internal control) signals are shown for the indicated groups. Scale bars, 100 μm. **c-d,** Maximum sequence identity distribution of 500 RNAverse-designed (**c**) gACA2b and (**d**) gACA36 variants. For each designed sequence, the maximum identity was calculated against all natural counterparts in the Rfam database. The blue dashed line indicates the maximum sequence identity of the rationally engineered RESTART construct to the same set of natural sequences. **e-f,** Functional evaluation of PTC-readthrough efficiency for 20 RNAverse-designed (**e**) gACA2b and (**f**) gACA36 variants in HEK293T cells. Relative eGFP intensity normalized to the Control group (top) and the relative fraction of eGFP-positive cells (bottom) were measured by flow cytometry. RNAverse-designed variants (yellow) are named and ordered by decreasing relative eGFP intensity. Data are presented as the mean **±** s.d from *n* = 3 biological replicates. The red dashed line indicates the mean performance of the RESTART construct. Asterisks indicate statistical significance for designs with higher mean values compared to the RESTART construct. *P*-values were determined using a two-sided Student’s *t*-test. \**P*≤0.05, \*\**P* ≤ 0.01, \*\*\**P* ≤ 0.001, \*\*\*\**P* ≤ 0.0001, n.s., not significant (*P* > 0.05).

**Extended Data Fig. 3.**
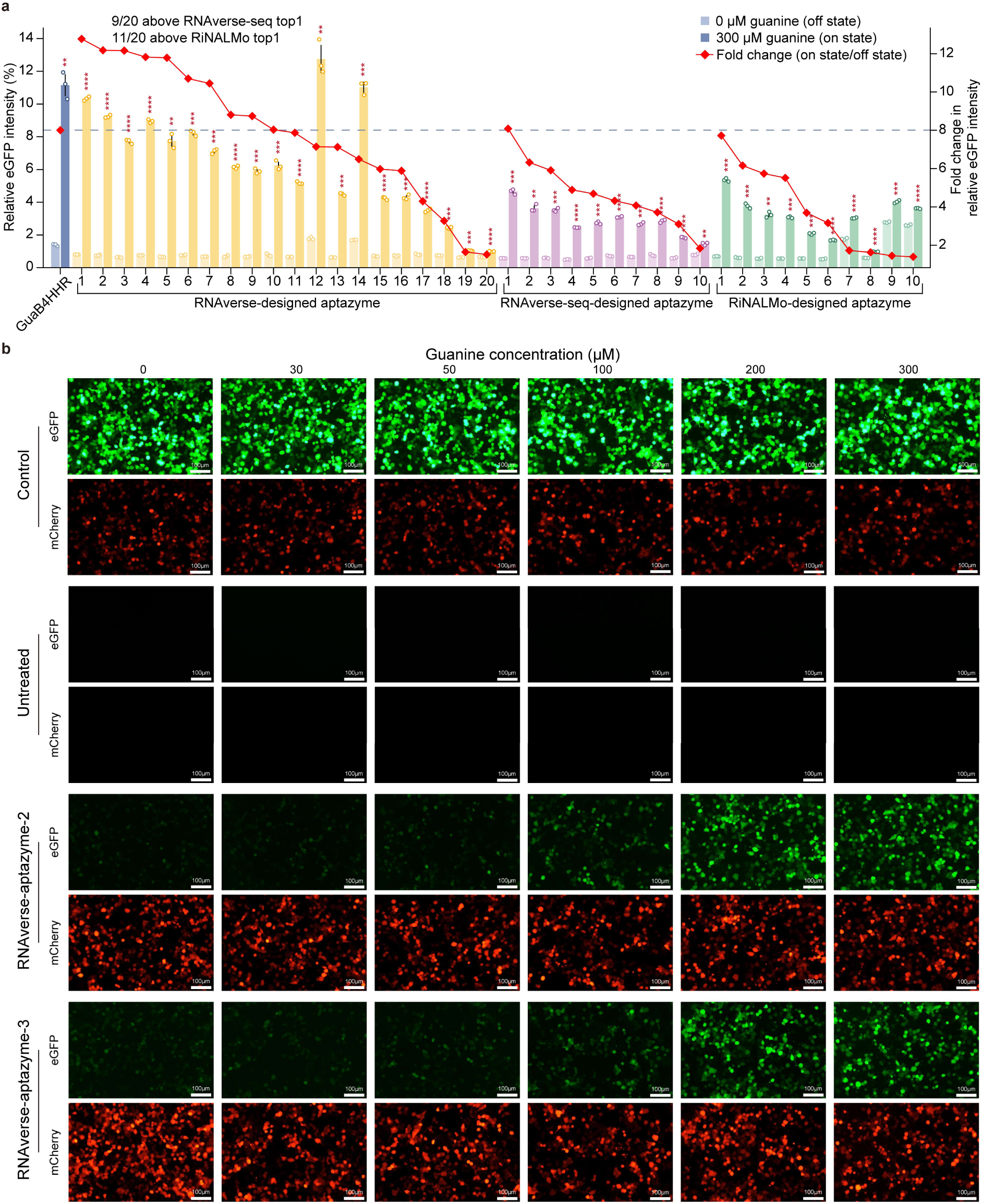
Supplemental data for aptazyme design and validation. **a,** Functional evaluation of designed guanine-responsive aptazymes in HEK293T cells. Relative eGFP intensity (left y axis) was measured for 20 RNAverse-designed (yellow), 10 RNAverse-seq-designed (purple), and 10 RiNALMo-designed (green) aptazymes, alongside the HTS-derived top variant GuaB4HHR (blue) by flow cytometry. For each design, light-colored bars represent the off state (0 μM guanine) and dark-colored bars represent the on state (300 μM guanine). The red line with diamonds indicates the fold change (on/off, right y axis) for each design. The blue dashed line indicates the fold change of GuaB4HHR. Variants within each group are named and ordered by decreasing fold change. Data are presented as the mean **±** s.d from *n* = 3 biological replicates. Asterisks indicate statistical significance between on and off states for each design. *P*-values were determined using a two-sided Student’s *t*-test. \**P*≤0.05, \*\**P* ≤ 0.01, \*\*\**P* ≤ 0.001, \*\*\*\**P* ≤ 0.0001; n.s., not significant (*P* > 0.05). RNAverse designs were compared against the top-performing design from each sequence-only baseline: 9/20 and 11/20 designs exceeded the top RNAverse-seq- and RiNALMo-designed variants in fold change, respectively. Bootstrap resampling (10 RNAverse designs sampled without replacement from the 20, 1,000 iterations) assessed robustness to unequal sample sizes; 74.4% and 95.3% of iterations were significant for RNAverse vs. RNAverse-seq and RNAverse vs. RiNALMo, respectively. Median numbers of RNAverse designs exceeding the baseline top1 in equal-sample bootstrap: 5/10 vs. RNAverse-seq and 6/10 vs. RiNALMo. **b,** Extended fluorescence images for aptazyme variants. Representative images of eGFP and mCherry expression under guanine concentrations ranging from 0 to 300 μM for Contro, Untreated, RNAverse-aptazyme-2, and RNAverse-aptazyme-3. Scale bar, 100 μm.

**Extended Data Fig. 4.**
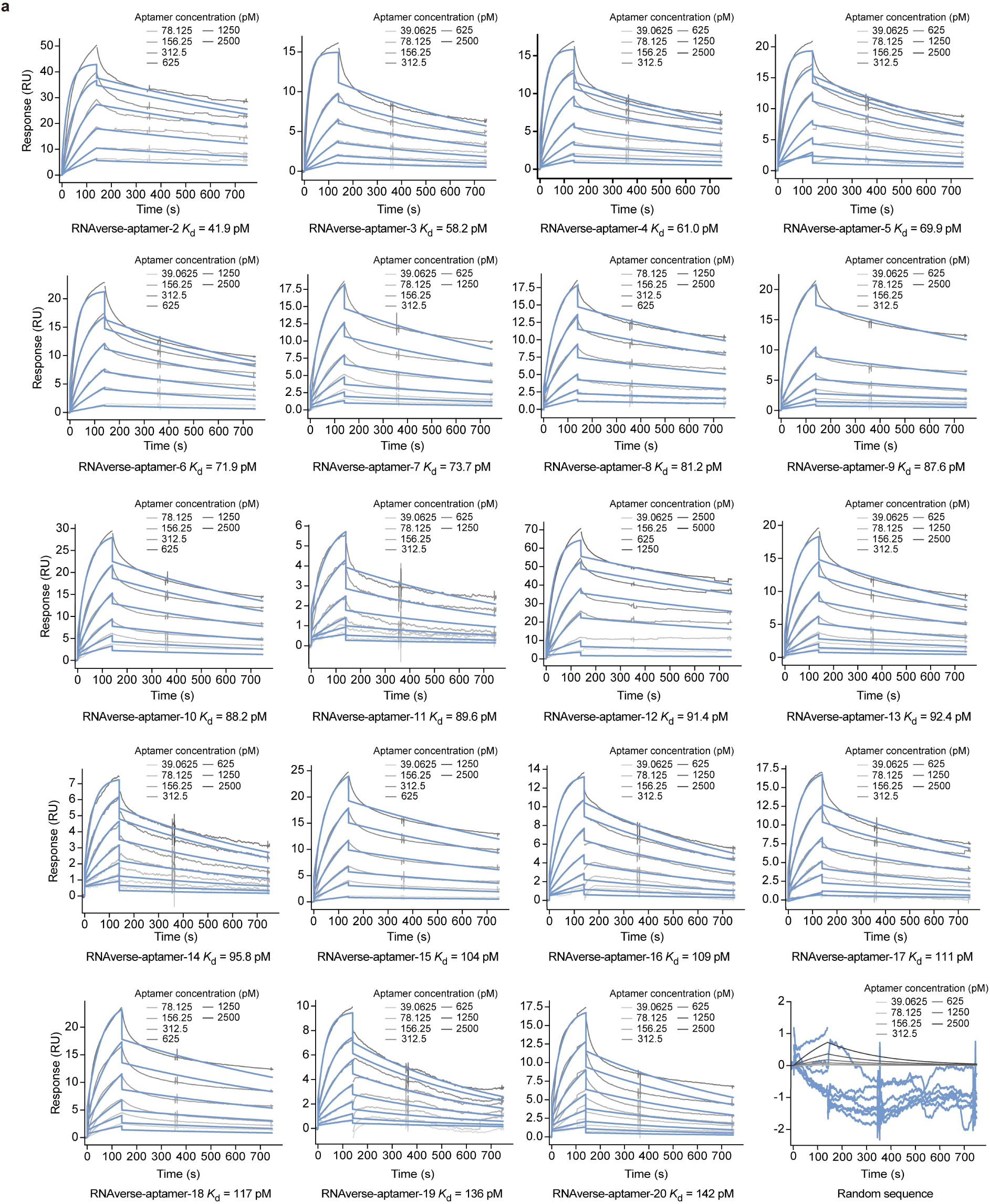
Binding kinetics of designed and benchmark IGFBP3 aptamers. **a,** SPR sensorgrams for the remaining 19 RNAverse-designed aptamers and a random negative control. For each variant, curves in varying shades of gray represent experimental binding responses (RU) at increasing analyte concentrations, and blue curves indicate the kinetic fits used for *K*_d_ calculation.

**Extended Data Fig. 5.**
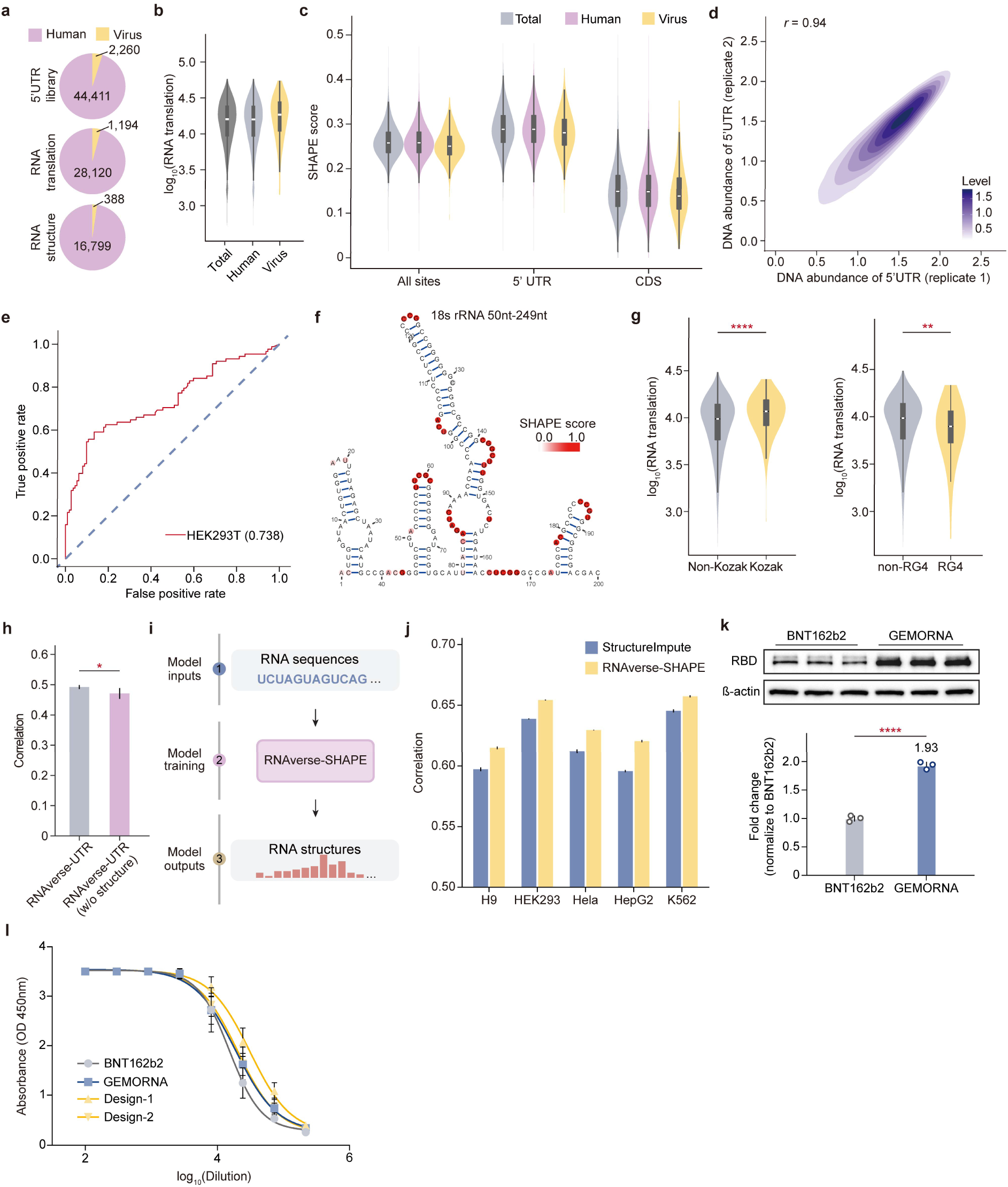
Dataset characterization, RNA structure profiling, and the RNAverse-guided 5′ UTR design pipeline. **a,** Composition of the high-throughput library, showing the proportion of human-(purple) and viral-derived (yellow) 5′ UTRs across the initial library, translation, and structure datasets. **b,** Distributions of RNA translation across the 5′ UTR library derived from human transcripts and viral genomes. **c,** Distributions of average SHAPE scores for the 150-nt 5′ UTR region (5′ UTR), the first 50-nt of the coding sequence (CDS), and the combined region (all sites) across the 5′ UTR library. **d,** Reproducibility of the profiling assay, showing Pearson correlations of DNA abundance of 5′ UTRs between biological replicates. **e,** Receiver operating characteristic (ROC) curve comparing SHAPE scores obtained from probing experiments with the reference 18S rRNA (50–249 nt region) secondary structure derived from the crystal structure. **f,** Reference secondary structure model of 18S rRNA (50–249 nt region). Nucleotides are colored by experimentally identified SHAPE scores. **g,** RNA translation of 5′ UTRs with or without canonical Kozak sequences (left) and RNA G-quadruplex (RG4) elements (right). *P* values were determined using a two-sided Student’s *t*-test. \**P*≤0.05, \*\**P* ≤ 0.01, \*\*\**P* ≤ 0.001, \*\*\*\**P* ≤ 0.0001; n.s., not significant (*P* > 0.05). **h,** Predictive performance of RNAverse-UTR compared with ablated variants without structural inputs. Correlation coefficients are shown. Data are presented as the mean **±** s.d over *n* = 5 cross-validation folds. *P*-values were determined using a two-sided paired Student’s *t*-test. \**P*≤0.05, \*\**P* ≤ 0.01, \*\*\**P* ≤ 0.001, \*\*\*\**P* ≤ 0.0001; n.s., not significant (*P* > 0.05). **i,** Schematic of the RNAverse-SHAPE framework for predicting RNA structure from sequence using training data collected across multiple cell lines. **j,** Predictive performance of RNAverse-SHAPE and StructureImpute. Correlation coefficients between predicted and experimentally identified SHAPE scores are shown for multiple cell lines. Data are presented as the mean **±** s.d from *n* = 3 replicates with different random seeds. **k,** Top, Representative western blot showing spike protein expression from BNT162b2 mRNA constructs bearing the original 5′ UTR and the 5′ UTR designed by GEMORNA. β-actin was used as a loading control. Bottom, Densitometric quantification of western blot signals. Band intensities were normalized to β-actin and then to the original BNT162b2 reference. Data are presented as mean ± s.d. from *n* = 3 biological replicates. *P*-values were determined using a two-sided Student’s *t*-test. \**P*≤0.05, \*\**P* ≤ 0.01, \*\*\**P* ≤ 0.001, \*\*\*\**P* ≤ 0.0001; n.s., not significant (*P* > 0.05). **l,** Serum antibody binding profiles and four-parameter logistic (4PL) curve fitting across vaccine formulations. Serial dilutions (log10-transformed) of serum collected from mice immunized with indicated vaccine formulations (BNT162b2, GEMORNA, RNAverse-UTR-designed 5′ UTRs; *n* = 5 mice per group) were evaluated for spike-specific IgG binding by ELISA. Solid curves represent non-linear regression using a 4PL model. Data are presented as mean ± s.d.

## Data availability

The model weights of RNAverse, RNAverse-aptamer, RNAverse-UTR, and RNAverse-SHAPE are available on figshare at https://doi.org/10.6084/m9.figshare.34476072. The benchmark datasets used in RNAverse are available on figshare at https://doi.org/10.6084/m9.figshare.34475931. Detailed Methods of RNAverse will be provided in the final publication.

## Code availability

The source code of RNAverse is available on GitHub at https://github.com/Chiyasa/RNAverse.

## Acknowledgements

The authors thank Q. Huang from the laboratory of Chengqi Yi for providing the snoRNA and DKC1-isoform3 constructs; and Y. Dong from the laboratory of Song-hai Shi for providing the AAV vector plasmid. Finally, we thank all members of the Q.W. laboratory and the Q.C.Z. laboratory for their valuable discussions and contributions. This work has been supported by the National Natural Science Foundation of China (Grants No. 32125007 and 32230018 to Q.C.Z.), the Science & Technology Fundamental Resources Investigation Program (Grants No. 2025FY100300 to Q.C.Z.), the Tsinghua University Dushi Program (Grants No. 20251080102 to Q.C.Z.), the TsienTang Life Science Development Fund at Tsinghua University (to Q.C.Z.), and the New Cornerstone Science Foundation through the XPLORER PRIZE (to Q.C.Z.).

## Author contributions

Q.C.Z. conceived the project. Q.C.Z. and Q.W. supervised the project. J.M. designed the RNAverse framework and conducted training. J.M., K.M., Z.D., R.G.H, and Y.W. collected and curated the pretraining and multimodal alignment datasets. J.M. collected downstream task datasets and performed model evaluation and benchmarking with the help of Y.S. K.M. and J.M. designed and implemented the in-painting algorithm. W.H. and J.M. designed and implemented the iterative optimization framework. J.Z. and J.M. performed the snoRNA and aptazyme experiments. K.H. performed the aptamer experiments. J.Z. and F.G. constructed the 5′ UTR library. W.H. designed the 5′ UTR library, selected sequences for library construction, and performed the bioinformatic analysis. J.Z. and K.H. conducted cellular validation experiments for the designed 5′ UTRs. X.X., K.H., J.Z., and L.Z. performed the mouse immunization experiments. Q.C.Z., J.M., J.Z., and W.H. wrote the manuscript with input from all authors.

